# Insulin-like peptides play distinct roles in nutrient-dependent plasticity in *Drosophila*

**DOI:** 10.1101/2025.08.12.669969

**Authors:** Michelle A. Henstridge, Bowen Slater, Jade R. Kannangara, Christen K. Mirth

## Abstract

The highly conserved insulin signalling pathway regulates growth and development time in response to nutrition across metazoans. The fruit fly, *Drosophila melanogaster* has eight insulin-like peptides, which are differentially expressed across development time, organs, and with nutritional conditions. However, whether individual insulin-like peptides play specific roles in controlling growth remains unknown. Recent studies have revealed that the ratio of protein to carbohydrates in the diet plays a key role in regulating life history traits, rather than the total caloric content of the diet alone. Furthermore, individual insulin-like peptides vary in their expression profiles according to nutrient conditions. Whether these differences in expression have any functional significance to animal life history traits remains unclear. Here we report that reducing the protein content of the larval diet through macronutrient restriction – where the calories lost from protein dilution are offset by increased carbohydrate content – results in a more pronounced developmental delay compared to caloric restriction – where both protein and carbohydrate concentrations are reduced. We further reveal that these two diet types result in notable differences in the expression levels of *Drosophila insulin-like peptides 2, 3* and *5*, and observe distinct phenotypic responses of individual *insulin-like peptide* mutants raised on each diet type. Taken together, our findings highlight the distinct roles of individual insulin-like peptides in regulating growth and development time in response to changes in dietary macronutrients, and provide key insights into the molecular mechanisms controlling nutritional plasticity in *Drosophila*.

**ARTICLE SUMMARY:** The insulin signaling pathway regulates growth and development in response to nutrition across metazoans. In the fruit fly, *Drosophila melanogaster,* seven insulin-like peptides have been identified that bind to a single insulin receptor, however their specific roles remain unclear. This study shows that reducing the protein content of the diet in different ways results in striking differences in the expression patterns of *insulin-like peptides* and different phenotypic responses of individual mutants raised on these diets. Together, these findings highlight the distinct roles insulin-like peptides play in regulating important life history traits in response to changes in dietary macronutrients.

## INTRODUCTION

The regulation of growth and development time in response to nutrition is mediated by the insulin/IGF-signalling (IIS) pathway, a highly conserved receptor-tyrosine kinase pathway found across all metazoans (Pertseva and Shpakov, 2002). Under high nutrition, insulin is secreted into circulation and binds to receptors on peripheral cells, initiating a signal transduction cascade resulting in the activation or repression of different genes that control growth (for review see Suzawa and Bland, 2023). Interestingly, many species are known to have multiple insulin-like peptides present in their genome (Pierce *et al*., 2001; Riehle *et al*., 2006; Gronke *et al*., 2010) and recent evidence suggests that these peptides may differ in both their expression patterns and biological functions (Semaniuk *et al*., 2020).

In *Drosophila,* eight insulin-like peptides (dILPs 1-8) have been identified (Brogiolo *et al*. 2001, Gronke *et al*., 2010, Garelli *et al*., 2012, Colombani *et al*., 2015), seven of which (dILPs1-7) are thought bind to a single insulin receptor (InR; Brogiolo *et al*., 2001; Garafolo, 2002). The various *dilp* genes are expressed in different larval and adult tissues and cell types (Brogiolo *et al*., 2001; Ikeya *et al*., 2002; Semaniuk *et al*., 2020), and therefore are likely to have different functions. For example, dILP6 is produced in the larval fat body and regulates growth during starvation and nonfeeding states (Okamoto *et al*., 2009; Slaidina *et al*., 2009; Bai *et al*., 2012), whereas dILPs 2, 3 and 5 are secreted by the insulin-producing cells (IPCs) within the brain and regulate growth in response to nutrition (Brogiolo *et al*., 2001; Rulifson *et al*., 2002; Geminard *et al*., 2009). Despite numerous studies on the roles of dILPs 2, 3 and 5 it is still not clear how these three peptides differ in their expression levels and function in response to changes in nutrition.

For many years research has focused on how total food intake influences development time and body size, rather than the specific composition of the food. Thus, diet manipulations have most often diluted all components of the diet, known as caloric restriction. In recent years, our understanding of the role of macronutrients in shaping the fitness of an organism has been greatly advanced by the development of nutritional geometry, a multidimensional modelling approach which enables the complex interactions between macronutrients, such as protein and carbohydrates, and life history traits to be examined (Lee et al., 2008; Simpson and Raubenheimer, 2012). Studies using nutritional geometry in *Drosophila* have revealed that traits like metabolism (Alton *et al.,* 2020), development time (Rodrigues *et al*., 2015; Kutz *et al*., 2019; Chakraborty *et al*., 2020; Kim *et al*., 2020; Min *et al*., 2021), body size (Rodrigues *et al*., 2015; Kutz *et al*., 2019; Chakraborty *et al*., 2020; Kim *et al*., 2020; Min *et al*., 2021), fecundity (Lee *et al*., 2008; Kim *et al*., 2020; Min *et al*., 2021), and lifespan (Lee *et al*., 2008; Skorupa *et al*., 2008; Kim *et al*., 2020; Min *et al*., 2021) are regulated more by specific ratios of protein to carbohydrates in the diet rather than total caloric content.

While the mechanism through which the protein to carbohydrate ratio of the diet affects these life history traits remains unknown, there are several lines of evidence to suggest that it involves changes in *dilp* expression (Kim and Neufeld, 2015; Post and Tatar, 2016; McDonald *et al*., 2021). In a recent study, McDonald *et al*. (2021) used a nutritional geometry framework to explore the effects of dietary protein and carbohydrate content on body size and the expression of genes that are transcriptionally regulated by the IIS pathway, including *dilps 2, 3, 5* and *8*. This research revealed dramatic differences in the expression patterns of each *dilp* across the nutritional landscape in female larvae, revealing a complex relationship between sex, diet and *dilp* expression (McDonald *et al*., 2021).

Here, we aimed to test whether the three brain-derived dILPs (dILPs 2, 3 and 5) are functionally different from one another by exploring whether they mediate different responses to diet in larval and adult life history traits. To do this, we compared the effects of macronutrient restriction, in which the calories lost from protein dilution are substituted by increasing the amount of carbohydrates, to caloric restriction, in which both protein and carbohydrate are diluted, on survival, development time, and body size. We then examined how larval expression levels of *dilps 2, 3* and *5* changed across protein concentrations and between diet types. Finally, we assessed how mutations in either *dilp2, 3* or *5* altered the phenotypic response to diet. Understanding the precise role of each dILP is critical to further our understanding of the mechanisms controlling nutrient-dependent plasticity in *Drosophila* and other organisms.

## MATERIALS AND METHODS

### Drosophila stocks and maintenance

The following stocks were used: *w^1118^* (BL5905), *w^1118^;TI{TI}ilp2^1^* (BL30881), *w^1118^;TI{TI}ilp3^1^*(BL30882), *w^1118^;TI{TI}ilp5^1^* (BL30884). All stocks were maintained at 25°C on a sucrose-yeast media containing, per litre: 100 g yeast (MP biomedicals), 50 g sugar (Bundaberg), 10 g agar (Gelita Australia), 30 mL Nipagen (10% in ethanol) and 3 mL propionic acid (Merck).

### Experimental diets

Experimental diets were based on manipulating the ingredients in a standard sucrose-yeast diet (Table 1; see recipe above), which contains a protein to carbohydrate (P:C) ratio of 1:2 and a caloric concentration of 495 calories/L. This control diet (1:2 or 100%) was then altered in two different ways: 1) caloric restriction, which reduced the caloric content of the diet to 50% and 25% of control food by reducing the amount of both sucrose and yeast while maintaining a constant P:C ratio (Table 1), or 2) macronutrient restriction, in which the calories from yeast were substituted by increasing the amount of sucrose, altering the P:C ratio to 1:5 and 1:10 while maintaining a constant number of calories (Table 1).

**Table 1.** Composition of experimental diets.

|  | *Amount (g/L)* | *Protein (g/L)* | *Carbs (g/L)* | *P+C* | *Calories* |
| --- | --- | --- | --- | --- | --- |
| *Control food (1:2 / 100%)* |  |  |  |  |  |
| Agar | 10 | 0 | 0 |  |  |
| Sugar | 50 | 0 | 50 |  |  |
| Yeast | 100 | 45 | 33 |  |  |
| Total |  | 45 | 83 | 128 | 495 |
| *Caloric restriction (50%)* |  |  |  |  |  |
| Agar | 10 | 0 | 0 |  |  |
| Sugar | 25 | 0 | 25 |  |  |
| Yeast | 50 | 22.5 | 16.5 |  |  |
| Total |  | 22.5 | 41.5 | 64 | 247.5 |
| *Caloric restriction (25%)* |  |  |  |  |  |
| Agar | 10 | 0 | 0 |  |  |
| Sugar | 12.5 | 0 | 12.5 |  |  |
| Yeast | 25 | 11.25 | 8.25 |  |  |
| Total |  | 11.25 | 20.75 | 32 | 123.75 |
| *Macronutrient restriction (1:5)* |  |  |  |  |  |
| Agar | 10 | 0 | 0 |  |  |
| Sugar | 89 | 0 | 89 |  |  |
| Yeast | 50 | 22.5 | 16.5 |  |  |
| Total |  | 22.5 | 105.5 | 128 | 495 |
| *Macronutrient restriction (1:10)* |  |  |  |  |  |
| Agar | 10 | 0 | 0 |  |  |
| Sugar | 108.5 | 0 | 108.5 |  |  |
| Yeast | 25 | 11.25 | 8.25 |  |  |
| Total |  | 11.25 | 116.75 | 128 | 495 |

### Survival, development time and body size analysis

Twenty-four hours after a 4 h lay on apple juice agar supplemented with yeast paste, first-instar larvae were placed into vials containing fly media (see recipe above) and scored every 8 h for the time taken to reach pupariation. At least 8-10 vials each containing 30 individual larvae for experiments using *w^1118^* or 10 larvae for *dilp* mutant/heterozygote experiments were collected for each genotype and diet. For measurements of pupal weight, individual pupae were removed from vials seven days after pupariation, washed with distilled water, dried and weighed on an XPR Ultra-Microbalance (Mettler Toledo). When measuring wing area, adult female flies were collected following eclosion and preserved in SH solution (70% ethanol, 30% glycerol) after their wings had fully expanded and hardened. The left wing was dissected off each body in SH solution under a Leica MZZ5 microscope, mounted onto microscope slides and digital images of the wings were captured using a Leica M165 FC stereoscope and a Leica DFC450 C digital camera. All wing images were analysed using ImageJ software, and wing area was calculated using landmarks from vein intercepts as per Pocas *et al*. 2022.

### Insulin-like peptide gene transcript quantification

To obtain newly molted third instar larvae for quantification of *dilp* expression levels, eggs were collected over a 4-hour laying interval on 55 millimeter diameter petri dishes containing each diet. Approximately 72-96 hours after egg lay (depending on diet type), larvae were carefully staged within 2 hours of the moult to the third larval instar as per Chakraborty *et al*. (2021). For each biological replicate, 10-15 third instar larvae (anterior end only) were snap frozen before RNA was extracted and DNAse treated using Direct-zol RNA MiniPrep kit (Zymo Research). Complementary DNA was synthesised using Tetro reverse transcriptase (Bioline) by priming 5µg of RNA with oligo-dT and random hexamers. Quantitative PCRs were performed in duplicate on a QuantStudio 3 (ThermoFisher) using SensiMix Sybr (Bioline) and primers specific for *dilp2* (F – 5’-ACG AGG TGC TGA GTA TGG TGT GCG-3’, R – 5’-CAC TTC GCA GCG GTT CCG ATA TCG-3’)*, dilp3* (F – 5’-GTC CAG GCC ACC AAT GAA GTT G-3’, R – 5’-CTT TCC AGC AGG GAA CGG TCT TCG-3’), *dilp5* (F – 5’-TGT TCG CCA AAC GAG GCA CCT TGG-3’, R – 5’-CAC GAT TTG CGG CAA CAG GAG TCG-3’) and *Rp49* (F – 5’-GCC GCT TCA AGG GAC AGT ATC T-3’, R – 5’-AAA CGC GGT TCT GCA TGA G-3’). Fold changes relative to *Rp49* were determined using the delta CT method and means and standard errors calculated from 3-5 biological replicates per genotype.

### Statistical analysis

Data was visualised and analysed in R (Rstudio version 1.4.1103). To visualise the data, we used the Tidyverse package (Wickham *et al*. 2019). Survival, development time and body size measurements were fitted with linear mixed effect models (lme4 package, Bates *et al*. 2015) using either survival, pupariation time or pupal volume/ wing area as response variables. The fixed effects included a second order polynomial for protein concentration, in addition to diet type, and genotype (where appropriate). Replicates were included as a random effect. Posthoc tests of the means and trends were conducted using the emmeans package (Lenth, 2025).

## RESULTS

To investigate how dietary composition effects larval viability, development time, and adult body size, we diluted the protein content of the larval diet in two ways: 1) caloric restriction, which reduces the caloric content of the diet to 50% and 25% of control food while maintaining a constant P:C ratio, or 2) macronutrient restriction, in which the calories from protein are substituted by increasing the amount of carbohydrates. This alters the P:C ratio from 1:2 in the control food to 1:5 and 1:10 while maintaining a constant caloric content. We then measured how raising larvae of a control genotype (*w^1118^*) on these different diets effected viability, development time, and body size.

On both the caloric and macronutrient restriction diets, egg to adult viability increased with increasing protein concentration (Fig. 1A, Table 2), however no significant differences in adult viability were found between the two diet types (Fig. 1A, Table 2). Development time was found to significantly decrease with increasing protein concentration on both the caloric and macronutrient restriction diets (Fig. 1B, Table 2). Interestingly, we observed a more severe developmental delay when larvae were raised on the low protein macronutrient restriction diets compared to the low protein caloric restriction diets (Fig. 1B, Table 2).

**Figure 1.**
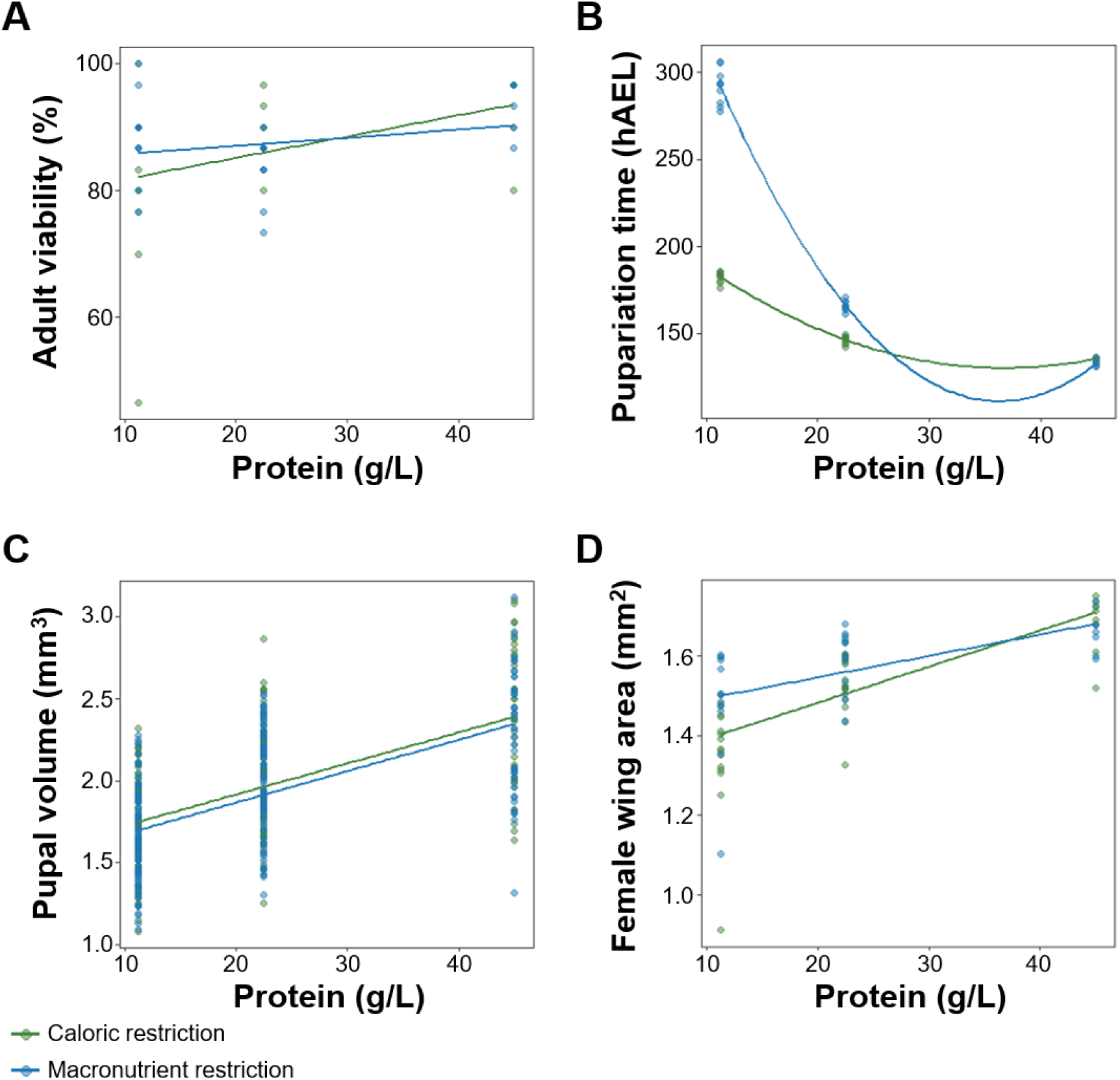
Dietary composition effects viability, development time and body size. **(A)** Adult viability increases with increasing protein concentration. No differences in viability were observed between larvae raised on the caloric restriction diet (green line) and the macronutrient restriction diet (blue line). Each data point represents one biological replicate of 30 animals. **(B)** Pupariation time decreases with increasing protein concentration, with a more profound effect in larvae raised on the macronutrient restriction diet. hAEL = hours after egg lay. **(C, D)** Final body size, measured by pupal volume **(C)** and female wing area **(D)**, increases with increasing protein concentration. Subtle differences in wing area were observed between the two diet types. For B-D, each data point represents a single animal. Alpha blending was applied to all graphs to indicate overlapping data points; darker areas represent higher point density.

**Table 2.** The effect of dietary manipulations on survival, pupariation time and body size.

| | Df | Viability<br>$\chi^2$ value | Pupariation time<br>$\chi^2$ value | Pupal Volume<br>$\chi^2$ value | Wing Area<br>$\chi^2$ value |
| --- | --- | --- | --- | --- | --- |
| Protein | 1 | 10.5045 ** | 91.525 *** | 209.707 *** | $2.574 \times 10^{-13}$ *** |
| Diet | 1 | 0.5874 | 42.177 *** | 2.2132 | 0.0411 * |
| Protein:Diet | 1 | 2.1628 | 27.104 *** | 0.0073 | 0.0652 |

Significance codes: \* $p < 0.05$ , \*\* $p < 0.01$ , \*\*\* $p < 0.001$

We found that final body size, as measured by pupal volume, increased with increasing protein concentration on both the caloric and macronutrient restriction diets (Fig. 1C, Table 2). However, despite the different effect diet type had on development time, we did not observe a significant difference of the impact of protein concentration on pupal volume between caloric and macronutrient restriction diets (Fig. 1C, Table 2). Like pupal volume, female wing area also increased with increasing protein concentration on both the caloric and macronutrient restriction diets (Fig. 1D, Table 2). However, subtle, but significant differences in female wing area were observed between the two diet types (Fig. 1D, Table 2). Together, these data suggest that diluting the protein content of the larval diet via macronutrient restriction has a different effect on larval growth and development time compared to protein dilution via caloric restriction.

Previous studies have revealed that dILPs secreted by the insulin producing cells differ in the way in which they respond to changes in dietary nutrient concentration (Kim and Neufeld, 2015; Post and Tatar, 2016; McDonald et al., 2021). Thus, the differences in development time we observed when *w^1118^* larvae were raised on the caloric and macronutrient restriction diets led us to hypothesise that these two methods of protein dilution may alter insulin signalling and, more specifically, *dilp* expression in different ways. To explore this idea, we measured *dilp2, 3* and *5* mRNA levels in third-instar *w^1118^*larvae that had been raised on either the caloric or macronutrient restriction diets.

As protein concentration increased, *dilp2* expression increased but only on the macronutrient restriction diet (Fig. 2A, Table 3). On the caloric restriction diet, we did not find a significant relationship between protein concentration and *dilp2* expression, suggesting that *dilp2* expression is repressed by high levels of carbohydrates but may not be affected by protein concentration. For *dilp3*, expression decreased with increasing protein concentration on both the caloric restriction and macronutrient restriction diets, with a steeper decrease in expression observed when protein was diluted via macronutrient restriction (Fig. 2B, Table 3). This suggests that *dilp3* expression is induced by high levels of carbohydrates and repressed by high levels of protein. Conversely, *dilp5* expression increased significantly with increasing protein concentration on both diet types, although this increase was more pronounced on the macronutrient restriction diet compared to the caloric restriction diet (Fig. 2C, Table 3). This expression pattern suggests that *dilp5* expression is induced by high levels of protein and repressed by high levels of carbohydrates. Taken together, these differing expression patterns suggest that the dILPs may function differently in response to changes in protein and carbohydrate concentration.

**Figure 2.**
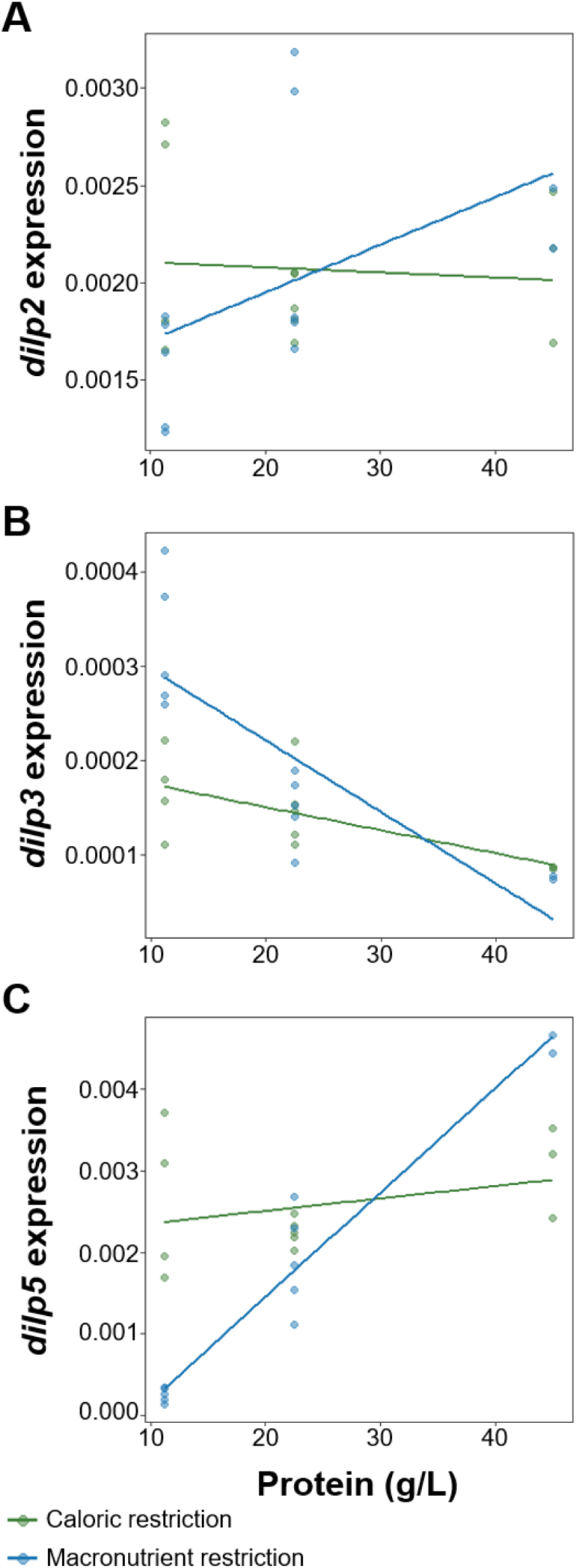
*dilp* expression patterns across diets. Diluting the protein content of the larval diet via caloric restriction (green line) or macronutrient restriction (blue line) alters the expression patterns of *dilp2* **(A)***, dilp3* **(B)**, and *dilp5* **(C)** differently. Expression levels were normalised to an internal control, *Rp49*. Each data point represents a biological replicate of 15 animals. Alpha blending was applied to indicate overlapping data points; darker areas represent higher point density.

**Table 3.** The effect of dietary manipulations on *dilp* expression levels.

| | Df | *dilp2* expression<br>$\chi^2$ value | *dilp3* expression<br>$\chi^2$ value | *dilp5* expression<br>$\chi^2$ value |
| --- | --- | --- | --- | --- |
| Protein | 1 | 1.8740 | 31.3344 *** | 53.720 *** |
| Diet | 1 | 0.0393 | 6.8208 ** | 10.734 ** |
| Protein:Diet | 1 | 4.4246 * | 9.2656 ** | 38.587 *** |

Significance codes: \* p<0.05, \*\* p<0.01, \*\*\* p<0.001

Given the striking differences we observed in the expression pattern of each *dilp*, both across protein concentrations and between diet types, we next investigated how individual null mutations in *dilps 2, 3* and *5* (Gronke *et al*., 2010) altered the development time and body size of larvae raised on either the caloric or macronutrient restriction diets. As observed for *w^1118^* (Fig. 1B, Table 2), the development time of all genotypes tested significantly decreased with increasing protein concentration on both diet types (Fig. 3, Table 4), and a more severe developmental delay was observed when larvae were raised on the low protein macronutrient diets (Fig. 3B, D, F, Table 4) compared to the low protein caloric restriction diets (Fig. 3A, C, E, Table 4). Final body size, as measured by female wing area, significantly increased with increasing protein concentration for all genotypes on both the caloric and macronutrient restriction diets (Fig. 4, Table 5), as seen for *w^1118^* (Fig. 1D, Table 2).

**Figure 3.**
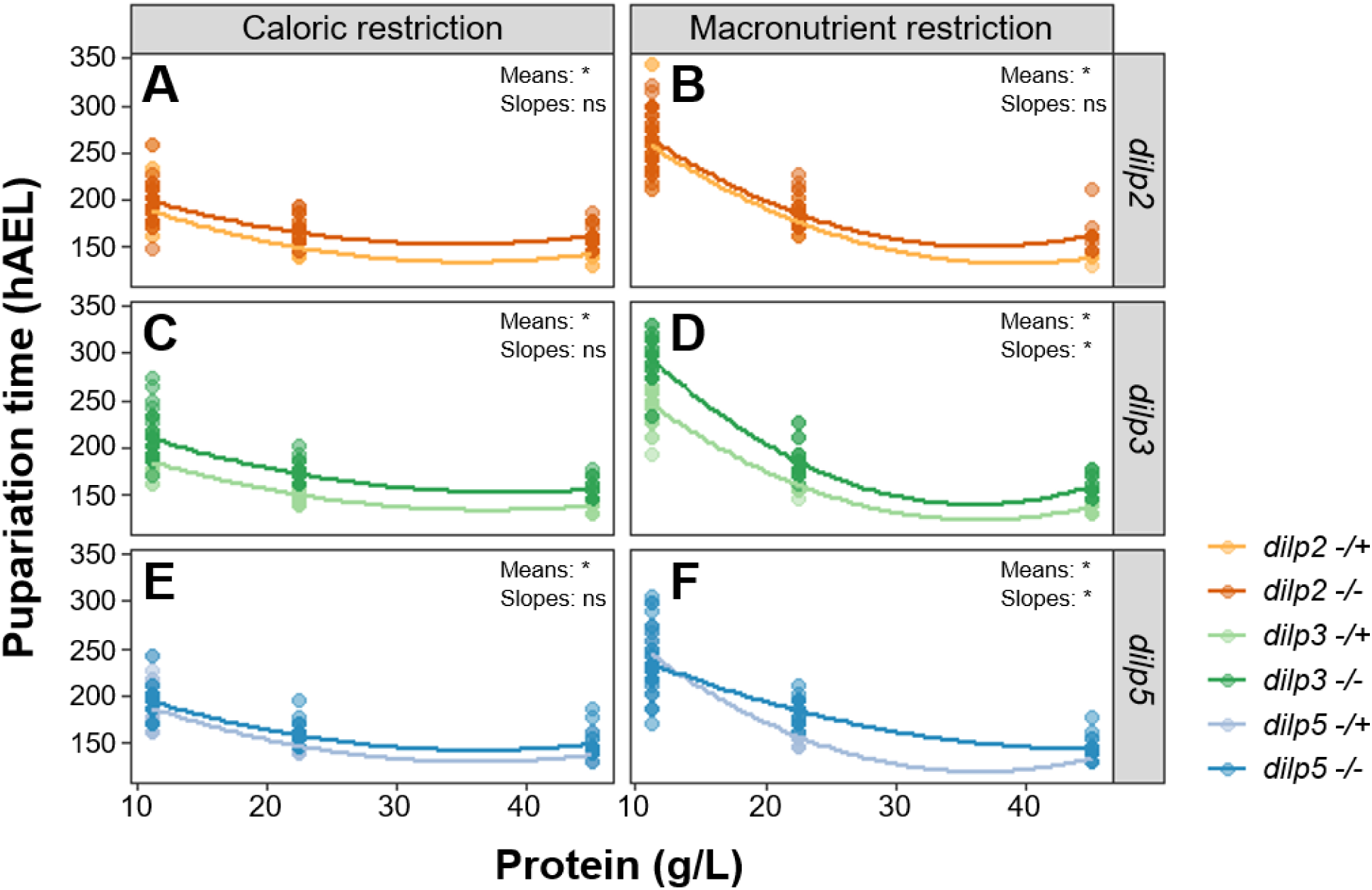
The developmental delay of *dilp* mutants is dependent on both dietary protein concentration and diet type. *dilp* mutants (-/-, darker lines) exhibit significantly longer mean pupariation times (means) compared to heterozygous controls (-/+, lighter lines) on both diet types **(A-F)**. For all genotypes, pupariation time decreases with increasing protein concentration **(A-F)**. The relationship between protein concentration and pupariation time (slopes) differs significantly between *dilp3* **(D)** and *dilp5* **(F)** mutants and their respective controls on the macronutrient restriction diet only; no significant differences in slopes were observed on the caloric restriction diet **(A, C, E)**. hAEL = hours after egg lay. Asterisks (*) indicate a significant difference (p < 0.05) in the mean or slope of pupariation time between mutants and controls; ns = not significant. Each data point represents a single animal. Alpha blending was applied to indicate overlapping data points; darker areas represent higher point density.

**Figure 4.**
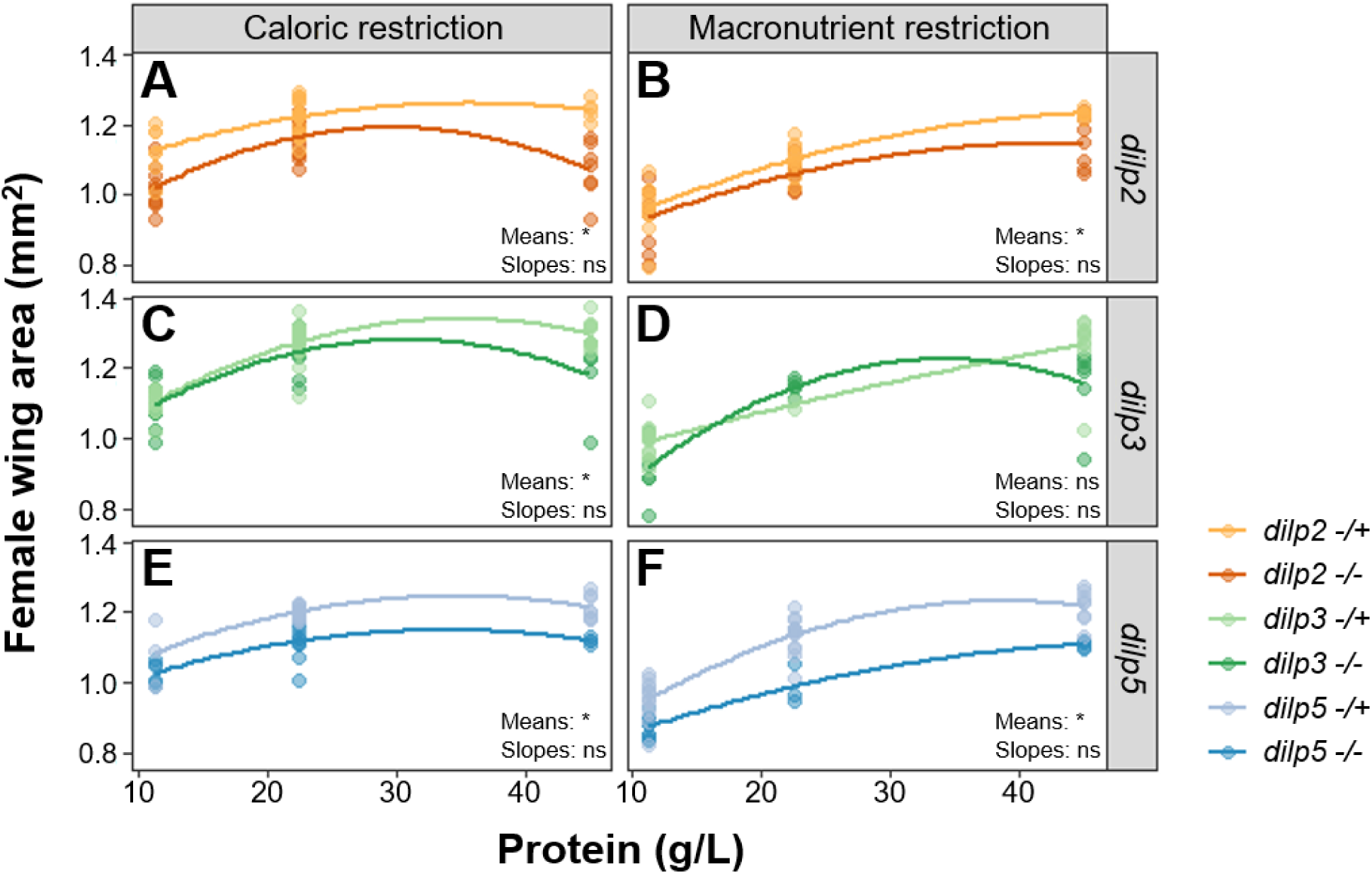
The relationship between dietary protein concentration and body size is not altered in *dilp* mutants, regardless of diet type. *dilp* mutants (-/-, darker lines) are significantly smaller than heterozygous controls (-/+, lighter lines) across both diet types (means), except for *dilp3* mutants on the macronutrient restriction diet **(D)**, where no significant difference in size is observed. For all genotypes, wing area increases with increasing protein concentration **(A-F)**. No significant differences in the relationship between protein concentration and wing area (slopes) were observed for any genotype on either diet type **(A-F)**. Asterisks (*) indicate a significant difference (p < 0.05) in the mean or slope of wing area between mutants and controls; ns = not significant. Each data point represents a single animal. Alpha blending was applied to indicate overlapping data points; darker areas represent higher point density.

**Table 4.** The effect of dietary manipulations on pupariation time in *dilp* mutants and heterozygotes.

| | Df | *dilp2* dev time<br>$\chi^2$ value | *dilp3* dev time<br>$\chi^2$ value | *dilp5* dev time<br>$\chi^2$ value |
| --- | --- | --- | --- | --- |
| Protein | 1 | 465.7739 *** | 640.0862 *** | 566.3273 *** |
| Diet | 1 | 239.3838 *** | 236.6184 *** | 148.0803 *** |
| Mutation | 1 | 20.6280 *** | 104.0495 *** | 12.3984 *** |
| Protein:Diet | 1 | 116.0699 *** | 118.8703 *** | 96.2492 *** |
| Protein:Mutation | 1 | 1.4665 | 5.2232 * | 7.5471 ** |
| Diet:Mutation | 1 | 1.7452 | 1.8269 | 0.4336 |
| Protein:Diet:Mutation | 1 | 0.0151 | 1.1902 | 2.1044 |

Significance codes: \* p<0.05, \*\* p<0.01, \*\*\* p<0.001

**Table 5.** The effect of dietary manipulations on female wing area in *dilp* mutants and heterozygotes.

| | Df | *dilp2* wing area<br>$\chi^2$ value | *dilp3* wing area<br>$\chi^2$ value | *dilp5* wing area<br>$\chi^2$ value |
| --- | --- | --- | --- | --- |
| Protein | 1 | 68.5664 *** | 98.3874 *** | 145.0874 *** |
| Diet | 1 | 51.6745 *** | 60.5688 *** | 83.0070 *** |
| Mutation | 1 | 37.0625 *** | 7.7068 ** | 76.7701 *** |
| Protein:Diet | 1 | 21.9174 *** | 9.3612 ** | 29.1894 *** |
| Protein:Mutation | 1 | 3.3742 | 3.9279 * | 0.1857 |
| Diet:Mutation | 1 | 3.2751 | 0.0024 | 1.6575 |
| Protein:Diet:Mutation | 1 | 0.0075 | 0.2500 | 0.6039 |

Significance codes: \* $p < 0.05$ , \*\* $p < 0.01$ , \*\*\* $p < 0.001$

When comparing the mean effect of individual *dilp* mutations to their heterozygous controls across all diets, we found that on both the caloric restriction and macronutrient restriction diets, *dilp2, dilp3* and *dilp5* mutants all had a significant developmental timing delay compared to their heterozygous controls (Fig. 3, Table 4). These delays did not always result in significant changes in body size. For example, averaging across all caloric restriction diets, *dilp2, dilp3* and *dilp5* mutants all had a significantly smaller wing area compared to their heterozygous controls (Fig. 4A, C, E, Table 5). However, when larvae were raised on the macronutrient restriction diets, only *dilp2* (Fig. 4B, Table 5) and *dilp5* (Fig. 4F, Table 5) mutants were significantly smaller than their corresponding heterozygous control. These data suggest that each dILP responds slightly differently to changes in protein concentration, and that this response differs depending on whether dietary protein is altered through caloric or macronutrient restriction.

Next, we analysed whether *dilp* mutant larvae responded differently to changes in protein concentration compared to heterozygous controls by comparing the slopes of the relationships between protein content in the diet and either development time or wing area. When larvae were raised on the caloric restriction diets no significant differences in the relationship between development time and protein concentration were observed for any of the genotypes (Fig. 3A, C, E). However, on the macronutrient restriction diets we found that *dilp3* (Fig. 3D) and *dilp5* (Fig. 3F) mutants responded differently to increasing protein concentration compared to their corresponding heterozygous controls. Specifically, we found that *dilp3* mutants had a steeper slope (Fig. 3D), indicative of a greater change in development time across protein concentrations, whereas *dilp5* mutants displayed a shallower slope (Fig. 3F), meaning that development time is less affected by protein concentration in this genotype.

For body size, no significant differences in the relationship between female wing area and protein concentration were observed between any of the genotypes on either the caloric or macronutrient restriction diets (Fig. 4). Thus, despite significant differences in the relationship between development time and protein concentration for some of the mutants analysed, this did not lead to changes in the relationship between protein concentration and body size. Taken together, these results demonstrate the important but distinct roles of dILPs 2, 3 and 5 in responding to dietary protein concentration and adjusting development and growth accordingly.

## DISCUSSION

Approaches using nutritional geometry have significantly advanced our understanding of the role macronutrients play in animal fitness (Lee *et al*., 2008; Simpson and Raubenheimer, 2012; Rodrigues *et al*., 2015; Alton *et al*., 2020). In *Drosophila*, nutritional geometry has revealed that the ratio of protein to carbohydrates in the diet is important for regulating life history traits, rather than the total caloric content alone (Lee *et al*., 2008; Skorupa *et al*., 2008). This was thought to be due to differences in *dilp* expression profiles across the nutritional landscape, however it was unclear if these differences were functionally significant (Kim and Neufeld, 2015; Post and Tatar, 2016; McDonald *et al*., 2021).

In this study, we diluted the protein content of the larval diet in two ways, caloric restriction and macronutrient restriction, and found striking differences in *dilp* expression patterns and development time both across protein concentrations and between diet types. Furthermore, our study revealed different growth phenotypes of individual *dilp* mutants raised on these diets. Taken together, our results support the hypothesis that changes in *dilp* expression are a key regulator of nutrient-dependent plasticity in *Drosophila* and suggest that each dILP may play slightly different roles in this process.

The observed expression patterns for *dilp3* and *dilp5* are consistent with previous studies which also found that *dilp3* expression declined with increasing protein concentration, whereas *dilp5* expression increased (McDonald *et al*., 2021; Post and Tatar, 2016). However, we note some inconsistencies in the expression pattern of *dilp2* observed in our study compared to others (McDonald *et al*., 2021; Post and Tatar, 2016). While we found that *dilp2* expression increased with increasing protein concentration on the macronutrient restriction diets, McDonald *et al*. (2021) have previously reported *dilp2* expression to decline with an increase in both protein and carbohydrates, which is also similar to the findings of Post and Tatar (2016). We did, however, find no significant change in *dilp2* expression across the caloric restriction diets, which has previously been shown (Ikeya *et al*., 2002; Geminard *et al*., 2009).

There are several possible explanations for these discrepancies. Firstly, our research measured *dilp* expression at the beginning of the third larval instar, whereas McDonald *et al*. (2021) took samples at the beginning of the wandering larval stage and Post and Tatar (2016) used adult flies in their study. As it is well known that IIS activity changes dynamically during development (Weger and Rittschof, 2024), and different *dilps* are expressed at different life stages (Brogiolo *et al*., 2001; Ikeya *et al*., 2002; Semaniuk *et al*., 2020), it is difficult to directly compare results of studies that have collected samples at different life stages. Secondly, while previous studies have used at least 24 different diets covering a broad nutrient space with total food concentrations ranging from 45 g/L to 360 g/L and P:C ratios ranging from 0:1 to 1.9:1 (Post and Tatar, 2016; McDonald *et al*., 2021), our study only included five different diets and thus covered a narrower nutrient space. The range of P:C ratios used in our study (1:10 – 1:2) fall within those previously used by Post and Tatar (2016) and McDonald *et* al (2021), whereas the total food concentrations of our diets (32 g/L – 128 g/L) goes below the lower limit of these previous studies, making direct comparisons between studies difficult. Future experiments should be designed such that *dilp* expression patterns can be measured in larvae raised across a broader range of diets to get a complete understanding of how macronutrients effect *dilp* expression levels.

Insulin/IGF-signalling is quite complex in *Drosophila*, with seven insulin-like peptides (dILPs 1-7) that are all thought to act through a single receptor, InR (Brogiolo *et al*., 2001; Garafolo, 2002). The extent of functional redundancy between dILPs, especially those with similar expression patterns, remains unclear. In a previous study, Gronke *et al*. (2010) showed that a combined *dilp2,3,5* gene-knockout had a more severe affect on development time, stress resistance and lifespan than individual *dilp* mutants, a finding which supports the idea that these peptides are redundant in their activation of the IIS pathway. However, our results suggest that this is not completely true. The striking differences in the expression patterns of *dilps 2, 3,* and *5* that were observed in response to changes in protein concentration, together with the diverse phenotypes of individual *dilp* mutants raised on these diets, provides strong evidence that dILPs 2, 3, and 5 have distinct roles in regulating growth and development time in response to changes in dietary macronutrients.

One important question that remains is how do the dILPs, which are thought to act through a single receptor, mediate a range of phenotypes in response to changes in protein concentration? The unique expression patterns of *dilps 2, 3* and *5* that we observed in response to changes in protein concentration could explain the specific phenotypes observed. However, if the dILPs are all binding to the same receptor why does the phenotype change depending on which individual dILP is bound? An alternative model is that dILPs differentially activate InR, possibly though different binding affinities, to induce distinct cell signalling outputs that result in specific downstream phenotypes. In favour of this model, a previous study has shown that two insulin-like peptide family members in the mosquito *Aedes aegypti* exhibit partially overlapping biological activity and different binding affinities for the mosquito InR, despite both being expressed in the brain (Wen *et al*., 2010). Similarly, it has been shown in *Drosophila* S2 cells that dILP2 and dILP5 differentially stimulate cell signalling and glycogen phosphorylase (Post *et al*., 2018). Future research should aim to test this model *in vivo* by examining the binding affinities of dILPs 2, 3 and 5 to InR.

Our study has provided important new insights into the distinct roles dILPs play in regulating nutrient-dependent plasticity in *Drosophila* that combines both expression data and important phenotypic information obtained from individual *dilp* mutants. Taken together, our results provide strong evidence to support the idea that the P:C ratio in the diet effects the expression of each *dilp* differently, and that these dILPs have differing, albeit overlapping, roles in the regulation of life history traits such as development time and body size. Future research should focus on a more in-depth analysis of both individual and combined *dilp* mutant animals on a wide nutritional landscape to tease apart the individual roles each dILP plays in regulating life history traits in response to changes in nutrition. Information gained from this research will greatly enhance our understanding of the mechanisms controlling nutrient-dependent plasticity in *Drosophila*.

## DATA AVAILABILITY

All strains are available upon request. We affirm that all data necessary for confirming conclusions of this article are present within the article, figures and tables. Datasets and R scripts are available at figshare: https://doi.org/10.26180/29826530.

### ACKNOWLEDGEMENTS

We thank Mia Wansbrough and the Australian *Drosophila* Biomedical Research Facility (OzDros) for technical support. M.A.H is a National Health and Medical Research Council (NHMRC) Early Career Fellow. This work was supported by an ARC Future Fellowship (FT170100259) and Discovery Project grant (DP240102830) to C.K.M.

## AUTHOR CONTRIBUTIONS

C.K.M conceived the experiments, interpreted the data and led the work. M.A.H conceived the experiments, performed the experiments and interpreted the data. B.S and J.R.K performed experiments. M.A.H and C.K.M wrote the manuscript with assistance from all authors.

## LITERATURE CITED

Alton, L. A., T. C. Kutz, C. L. Bywater, J. E. Beaman, P. A. Arnold et al., 2020 Developmental nutrition modulates metabolic responses to projected climate change. Funct. Ecol. 34: 2488–2502.

Bai, H., P. Kang, and M. Tatar, 2012 *Drosophila* insulin-like peptide-6 (dilp6) expression from fat body extends lifespan and represses secretion of *Drosophila* insulin-like peptide-2 from the brain. Aging Cell 11: 976–985.

Bates, D., M. Machler, B. Bolker, and S. Walker, 2015 Fitting linear mixed-effects models using lme4. J. Stat. Softw. 67: 1–48.

Brogiolo, W., H. Stocker, T. Ikeya, F. Rintelen, R. Fernandez et al., 2001 An evolutionarily conserved function of the *Drosophila* insulin receptor and insulin-like peptides in growth control. Curr. Biol. 11: 213–221.

Chakraborty, A., C. M. Sgro, and C. K. Mirth, 2020 Does local adaptation along a latitudinal cline shape plastic responses to combined thermal and nutritional stress? Evol. 74:2073–2087.

Chakraborty, A., C. M. Sgro, and C. K. Mirth, 2021 The proximate sources of genetic variation in body size plasticity: The relative contributions of feeding behaviour and development in *Drosophila melanogaster*. J. Insect Physiol. 135: 104321.

Colombani, J., D. S. Andersen, L. Boulan, E. Boone, N. Romero et al., 2015 *Drosophila* Lgr3 couples growth with maturation and ensures developmental stability. Curr. Biol. 25: 2723–2729.

Garafolo, R. S., 2002 Genetic analysis of insulin signalling in *Drosophila*. Trends Endocrinol. Metab. 13: 156–162.

Geminard, C., E. J. Rulifson, and P. Leopold, 2009 Remote control of insulin secretion by fat cells in *Drosophila*. Cell Metab. 10: 199–207.

Grönke, S., D. F. Clarke, S. Broughton, T. D. Andrews, and L. Partridge, 2010 Molecular evolution and functional characterization of *Drosophila* Insulin-like peptides. PLoS Genet. 6: e1000857.

Ikeya, T., M. Galic, P. Belawat, K. Nairz, and E. Hafen, 2002 Nutrient-dependent expression of insulin-like peptides from neuroendocrine cells in the CNS contributes to growth regulation in *Drosophila*. Curr. Biol. 12: 1293–1300.

Kim, J., and T. P. Neufeld, 2015 Dietary sugar promotes systemic TOR activation in *Drosophila* through AKH-dependent selective secretion of Dilp3. Nat. Commun. 6: 6846.

Kim, K. E., T. Jang, and K. P. Lee, 2020 Combined effects of temperature and macronutrient balance on life-history traits in *Drosophila melanogaster:* implications for life-history trade-offs and fundamental niche. Oecologia 193: 299–309.

Kutz, T. C., C. M. Sgro, and C. K. Mirth, 2019 Interacting with change: Diet mediates how larvae respond to their thermal environment. Funct. Ecol. 33:1940–1951.

Lee, K. P., S. J. Simpson, F. J. Clissold, R. Brooks, J. W. Ballard et al., 2008 Lifespan and reproduction in *Drosophila*: New insights from nutritional geometry. Proc. Natl. Acad. Sci. 105: 2498–2503.

Lenth, R., 2025 emmeans: Estimated marginal means, aka least-squares means. R package version 1.11.

McDonald, J. M., P. Nabili, L. Thorsen, S. Jeon, and A. W. Shingleton, 2021 Sex-specific plasticity and the nutritional geometry of insulin-signaling gene expression in *Drosophila melanogaster*. EvoDevo 12: 6.

Min, K. W, T. Jang, and K. P. Lee, 2021 Thermal and nutritional environments during development exert different effects on adult reproductive success in *Drosophila melanogaster*. Ecol. Evol. 11: 443–457.

Okamoto, N., N. Yamanaka, Y. Yagi, Y. Nilshida, H. Kataoka et al., 2009 A fat body-derived IGF-like peptide regulates postfeeding growth in *Drosophila*. Dev. Cell 17: 885–891.

Pertseva, M. N., and A. O. Shpakov, 2002 Conservatism of the insulin signaling system in evolution of intertebrate and vertebrate animals. J. Evol. Biochem. Phys. 38: 547–561.

Pierce, S. B., M. Costa, R. Wisotzkey, S. Devadhar, S. A. Homburger et al., 2001 Regulation of DAF-2 receptor signaling by human insulin and ins-1, a member of the unusually large and diverse *C. elegans* insulin gene family. Genes Dev. 15: 672–686.

Pocas, G. M., A. E. Crosbie, and C. K. Mirth, 2022 When does diet matter? The roles of larval and adult nutrition in regulating adult size traits in *Drosophila melanogaster*. J. Insect Physiol. 139: 104051.

Post, S., and M. Tatar, 2016 Nutritional geometric profiles of Insulin/IGF expression in *Drosophila melanogaster*. PLoS One 11: e0155628.

Post, S., G. Karashchuk, J. D. Wade, W. Sajid, P. De Meyts et al., 2018 *Drosophila* insulin-like peptides DILP2 and DILP5 differentially stimulate cell signalling and glycogen phosphorylase to regulate longevity. Front. Endocrinol. 9: 245.

Riehle, M. A., Y. Fan, C. Cao, and M. R. Brown, 2006 Molecular characterisation of insulin-like peptides in the yellow fever mosquito, *Aedes aegypti*: Expression, cellular localisation, and phylogeny. Peptides 27: 2547–2560

Rodrigues, M. A., N. E. Martins, L. F. Balance, L. N. Broom, A. J. S. Dias et al., 2015 *Drosophila melanogaster* larvae make nutritional choices that minimize developmental time. J. Insect Physiol. 81: 69–80.

Rulifson, E. J., S. K. Kim, and R. Nusse, 2002 Ablation of insulin-producing neurons in flies: growth and diabetic phenotypes. Science 296: 1118–1120.

Scorupa, D. A., A. Dervisefendic, J. Zwiener, and S. D. Pletcher, 2008 Dietary composition specifies consumption, obesity, and lifespan in *Drosophila melanogaster*. Aging Cell 7: 478–490.

Semaniuk, U., V. Piskovatska, O. Strilbytska, T. Strutynska, N. Burdyliuk et al., 2020 *Drosophila* insulin-like peptides: from expression to functions – a review. Entom. Exp. Appl. 169: 195–208.

Slaidina, M., R. Delanoue, S. Gronke, L. Partridge, and P. Leopold, 2009 A *Drosophila* insulin-like peptide promotes growth during nonfeeding states. Dev. Cell 17: 874–884.

Suzawa, M., and M. L. Bland, 2023 Insulin signaling in development. Dev. 150: dev201599.

Weger, A. A., and C. C. Rittschof, 2024 The diverse roles of insulin signalling in insect behaviour. Front. Insect Sci. 4: 1360320.

Wen, Z., M. Gulia, K. D. Clark, A. Dhara, J. W. Crim et al., 2010 Two insulin-like peptide family members from the mosquito *Aedes aegypti* exhibit differential biological and receptor binding activities. Mol. Cell Endocrinol. 328: 47–55.

Wickham, H., M. Averick, J. Bryan, W. Chang, L. D. McGowan et al., 2019 Welcome to the tidyverse. J. Open Source Softw. 4: 1686.

